# Enhancing genome recovery across metagenomic samples using MAGmax

**DOI:** 10.1101/2025.05.28.656617

**Authors:** Arangasamy Yazhini, Johannes Söding

## Abstract

**Summary:** The number of metagenome-assembled genomes (MAGs) is rapidly increasing with the growing scale of metagenomic studies, driving fast progress in microbiome research. Sample-wise assembly has become the standard due to its computational efficiency and strain-level resolution. It requires dereplication, the removal of near-identical genomes assembled in different metagenomic samples. We present MAGmax, an efficient dereplication tool that enhances both the quantity and quality of MAGs through a strategy of bin merging and re-assembly. Unlike dRep, which selects a single representative bin per genome cluster, MAGmax merges multiple bins within a cluster and reassembles them to increase coverage. MAGmax produces more dereplicated, higher-quality MAGs than dRep at 1.6*×* its speed and using three times less memory.

**Availability and implementation:** The MAGmax open source software, implemented in Rust, is available under the GPLv3 license at https://github.com/soedinglab/MAGmax.

## 1 Introduction

Shotgun sequencing of environmental samples has enabled the recovery of metagenome-assembled genomes (MAGs), providing insights into the ecology and function of uncultured microbes. The standard computational workflow for reconstructing MAGs consists of the following steps: assembly of metagenomic reads, binning of assembled contigs according to their genome of origin, and dereplication of bins [1]. In large-scale studies, each sample is typically processed independently for assembly and binning [2]. This sample-wise strategy offers greater computational efficiency compared to co-assembly across all samples, and is particularly advantageous for reconstructing strain-level genomes, as it reduces the complexity of the deBruijn graph during assembly [3].

Since the same genomes may be present in multiple metagenomic samples, bins obtained from sample-wise assemblies are often redundant. Redundancy is typically addressed by clustering bins that belong to the same genome based on average nucleotide identity (ANI) and selecting a representative bin per cluster based on quality metrics such as genome completeness and purity. A widely used tool for this dereplication step is dRep [4], which uses MASH to precluster bins and then applies FastANI [5] or skani [6] to define more refined secondary clusters. For each secondary cluster, dRep selects the highest-quality bin as the representative, using CheckM1 to estimate completeness and contamination [7].

The current dereplication approach has several limitations: (i) It discards bins with completeness below the predefined threshold, even if they are highly pure, throwing away valuable genomic information. (ii) MASH underestimates ANI for bins with less than 90% completeness [6], which can result in grouping incomplete bins from the same genome into separate primary clusters. (iii) The average linkage clustering algorithm used by dRep may fail to cluster together bins with ANI above the threshold. (iv) Even when ANI estimates are accurate and bins are correctly grouped, only a single representative bin is retained while other bins are discarded that may contain genomic regions not covered by the selected bin [8]. (v) dRep does not support CheckM2, which provides more accurate estimates of completeness and contamination than CheckM1 [9].

We introduce MAGmax, a tool that improves the yield and quality of metagenome-assembled genomes (MAGs) through bin merging and reassembly, addressing limitations of dRep. MAGmax integrates sequences from all bins in a cluster, including those with low completeness but high purity, and reassembles them to enhance bin quality. It uses skani, a state-of-the-art method for accurate ANI estimation, and CheckM2 for assessing bin quality.

## 2 Results

### Algorithm overview

MAGmax takes sample-specific genomic bins as input and filters them based on a user-defined purity threshold (default: *>* 95%). It identifies single-linkage connected components among these bins based on average nucleotide identities (ANI, default: 99%) between bin pairs, using skani [6] and a depth-first search algorithm. Within each component, clusters are formed using maximal clique detection. In this approach, a bin can belong to multiple clusters. The algorithm ensures that all bin pairs with sequence identity above the ANI cutoff are grouped together. Bins that are not part of any cluster are added to an existing cluster if they share ANI above the cutoff only with at least one cluster member. For each cluster, MAGmax selects the representative bin with the highest quality score, defined as completeness – 5 *×* contamination, where completeness must be *>* 90% and contamination *<* 5%. If no such bin exists, the bins within the cluster are merged and reassembled using SPAdes [10]. The quality score of the reassembled bin is then compared with that of original input bins and the bin with the best quality score is selected. Finally, MAGmax performs a round of redundancy removal, retaining only the best-quality bin from any pair sharing an ANI above the cutoff (Fig. **1**a).

### Output

A non-redundant set of dereplicated genomic bins, including bins improved through merging and reassembly, and a text file listing bin completeness and contamination estimated by CheckM2 [9].

We evaluated the performance of MAGmax on metagenomic samples from three different datasets: 15 gut samples from villagers in Honduras [11], 18 samples from black soil [12] and 43 gut samples from newborns at day 21 postpartum [13]. To obtain genomic bins, we binned sample-wise assembled contigs with multi-sample coverage using GenomeFace [14], VAMB [15] and MetaBAT2 [16] (Materials). MAGmax and dRep were independently applied to dereplicate bins at the strain-level (ANI 99%).

First, we compared the number of bins produced by dRep and MAGmax as a function of quality score, using bins with ≥ 50% completeness and *<* 5% or *<* 10% contamination. In the dRep output, some bin pairs remained redundant with ANI above 99% (Supplementary Table 1). To avoid redundancy in all subsequent analyses, we removed the lower-quality bin within each redundant pair. Results presented in the main text correspond to bins with contamination *<* 5%.

The total number of de-replicated bins obtained using MAGmax is consistently higher than those produced by dRep (Fig. 1b). In the GenomeFace results, which generated the highest number of bins among the three binning tools, MAGmax recovered 37, 1, and 6 more bins than dRep for the human gut (Honduras), black soil, and neonatal gut samples, respectively. The largest gain was seen in the MetaBAT2 results, where MAGmax yielded 52, 2, and 18 additional bins across the same datasets. Similar results were observed for contamination level *<* 10% (Supplementary Fig. 1).

**Figure 1.**
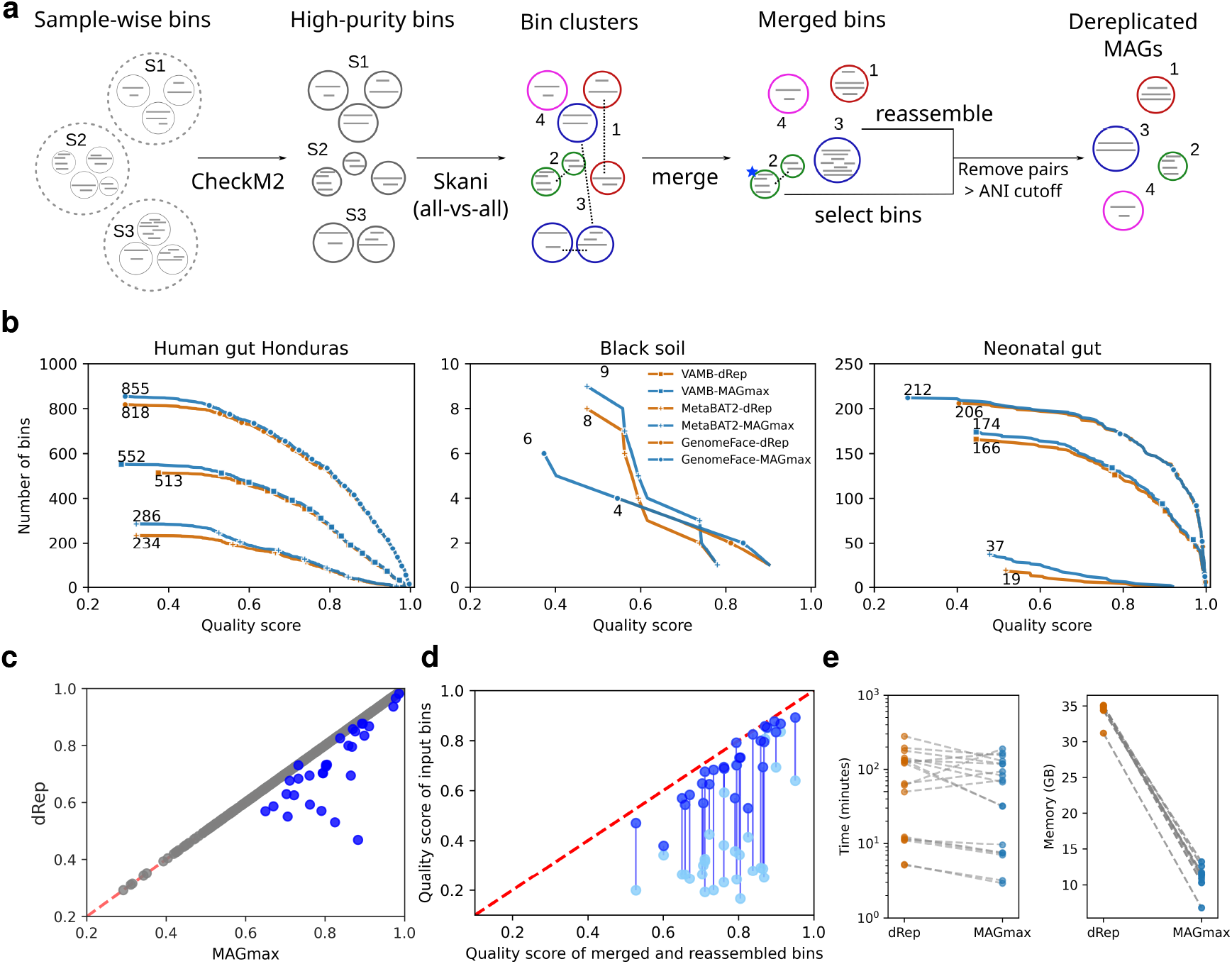
**a**, Overview of MAGmax. The blue star marks a high-quality bin. **b**, Complementary cumulative distribution of the number of dereplicated genomic bins over the bin quality score (completeness – 5 *×* purity) in combination with three popular binners. **c**, Quality scores of dRep bins (y-axis) and the corresponding MAGmax bins (x-axis) based on genomic cluster membership. Grey/blue circles: MAGmax and dRep bins are identical/different. **d**, Scatter plot showing quality scores of merged and reassembled bins (x-axis) versus input bins (y-axis). Blue circles represent the highest-quality bins within each genomic cluster, while cyan circles indicate the other input bins. Vertical lines connect the input bins that were used for merging and reassembly. **e**, Runtime (in minutes) and peak memory usage (in GB).

Next, we compared the quality of each dRep bin with the corresponding output bin from MAGmax, which could be the same bin, a different representative bin from its genomic cluster, or a merged and reassembled bin. In Fig. **1**c, GenomeFace bins from the Honduras gut dataset showed that 96.1% of bins were the same between two methods (grey circles), while 3.9% differed, with MAGmax bins having consistently higher quality than dRep bins (blue circles). Across the binning tools, an average of 95% of bins were selected commonly by both methods and the remaining 5% consistently showed higher quality scores in MAGmax (Supplementary Fig. 2a). This trend was consistent across the Honduras gut and neonatal gut datasets.

In the black soil dataset, VAMB produced only 10 bins, from which MAGmax produced one non-redundant bin. The dRep run failed as no clusters were formed during the primary clustering step with MASH [17]. For MetaBAT2, dRep and MAGmax selected identical bins. For GenomeFace, MAGmax produced one bin that differed from dRep and had a higher quality score (Supplementary Fig. 2a). Results for 10% contamination were consistent with those for 5% (Supplementary Fig. 2b).

Since dRep uses CheckM1 to select representative bins, we evaluated MAGmax’s performance based on CheckM1 quality scores (Supplementary Fig. 2). Bins from MAGmax that differ from dRep within the same genomic cluster consistently show higher quality according to CheckM1 predictions, indicating that MAGmax produces superior bins compared to those selected by dRep.

To assess the impact of merging and reassembly, we compared the quality scores of merged and reassembled out bins with the corresponding input bins. Fig. **1**d shows example results from GenomeFace bins for the Honduras gut dataset, where output bins exhibited substantial improvement in the quality, with scores increasing by 0.2% to 36.9% compared to the highest quality input bin (dark blue). For instance, two input bins with quality scores *<* 0.4 were merged and reassembled into a single bin with a score of 0.6, demonstrating the effectiveness of this strategy. Similar trends were observed across other binning tools, datasets, and at the 10% contamination level (Supplementary Fig. 3).

Across all datasets and binning tools, MAGmax was on average 1.6 *×* faster than dRep, while using one-third of the peak memory (Fig. **1**e). The gain in speed is largely attributed to MAGmax’s integration of CheckM2, which is considerably faster than CheckM1 used by dRep [9].

In conclusion, MAGmax enhances both the quality and quantity of metagenome-assembled genomes (MAGs) through dereplication and bin enrichment. It allows users to set desired cutoffs for ANI, completeness, and contamination. With the exponential increase in metagenomic data, MAGmax’s ability to integrate genomic information across multiple samples, coupled with its speed and memory efficiency, will improve the recovery of MAGs in large-scale metagenomic studies.

## Methods

### Datasets preprocessing

For benchmarking, we selected 76 real metagenomic samples from three different environments: 15 human gut samples from villagers in Honduras (BioProject accession: PRJNA999635) [11], 18 from black soil (BioProject accession: PRJNA1226397) [12], and 43 from neonatal gut (Study Accession: ERP115334) [13]. Raw reads from black soil were error-corrected using Musket (v1.1) [18], while reads from gut samples were processed using kneadData (v0.12.0) to perform error correction and remove human DNA contamination using the GRCh37 reference assembly. Corrected reads were sample-wise assembled using SPAdes (v4.0.0) [10]. For neonatal samples, MEGAHIT (v1.2.9) was used for assembly. Read mapping was performed using Strobealign (v0.13.0-25-g3a97f6b) [19] to generate abundance matrices and alignment files, followed by sorting using samtools (v1.19) [20].

### Binning

Contigs from all samples within the same environment were concatenated, and binning was performed using GenomeFace [14], VAMB [15], and MetaBAT2 [16] with multi-sample coverage. For GenomeFace and VAMB, contigs longer than 1 kb were used, while MetaBAT2 required contigs longer than 1.5 kb. Bins generated by MetaBAT2 were manually split by sample IDs, while GenomeFace and VAMB produced sample-wise bins directly. Only bins with a total sequence length of at least 200 kb were considered for dereplication.

### Benchmarking runs

Bins were dereplicated at the strain-level (99% ANI) using dRep [4] with the options `--S algorithm skani, -sa 0.95, -pa 0.99, -comp 50, -p 24` and `-cont 5 or -cont 10`. MAGmax was run on the same input bins with the options `-c 50, --threads 24 and -p 95 or -p 90`. Runtime and peak memory usage were evaluated on a Linux cluster system using 1 CPU with 24 cores and 128 GB of RAM.

### Bin quality assessment

CheckM2 [9] was used to estimate bin completeness and contamination. The quality score was calculated as completeness - 5 *×* contamination [21], and compared between bins obtained from dRep and MAGmax. dRep’s de-replicated bins were further filtered based on completeness and purity estimated by CheckM2. MAGmax output was assessed and filtered using completeness and contamination values estimated using CheckM1 with the default settings [7].

## Funding

YA acknowledges support from Marie Sklodowska-Curie Actions (Project No. 101111457) under the Horizon Europe programme of the European Union and from the Max-Planck society.

## Conflict of Interest

none declared.

## Notes

### Competing Interest Statement

The authors have declared no competing interest.

