## Supplementary Materials for "Enhancing genome recovery across metagenomic samples using MAGmax"

### Supplementary Tables

Table 1: The number of bin pairs with ≥99% ANI in the dRep output

| Binner | Dataset | Contamination level (%) | No. of pairs |
|---|---|---|---|
| VAMB | Human gut Honduras | 5 | 1 |
| MetaBAT2 | Human gut Honduras | 5 | 1 |
| GenomeFace | Human gut Honduras | 5 | 2 |
| VAMB | Neonatal gut | 5 | 2 |
| GenomeFace | Neonatal gut | 5 | 6 |
| VAMB | Human gut Honduras | 10 | 1 |
| MetaBAT2 | Human gut Honduras | 10 | 1 |
| GenomeFace | Human gut Honduras | 10 | 2 |
| VAMB | Black soil | 10 | 1 |
| VAMB | Neonatal gut | 10 | 3 |
| GenomeFace | Neonatal gut | 10 | 4 |

### Supplementary Figures

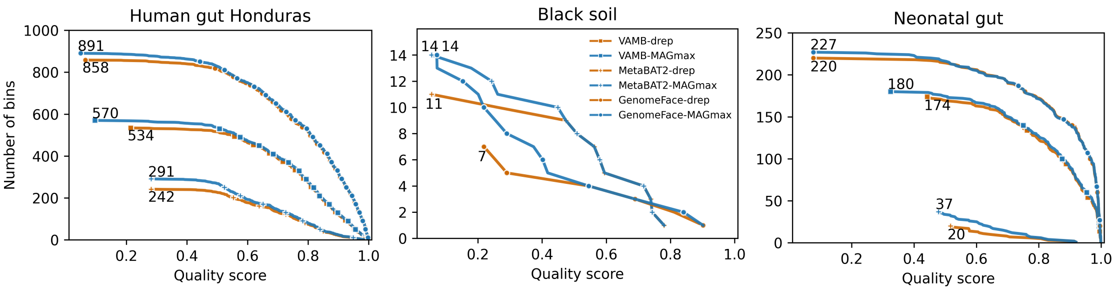


**Figure 1**: Complementary cumulative frequency of dereplicated bins as a function of quality score at 10% contamination.

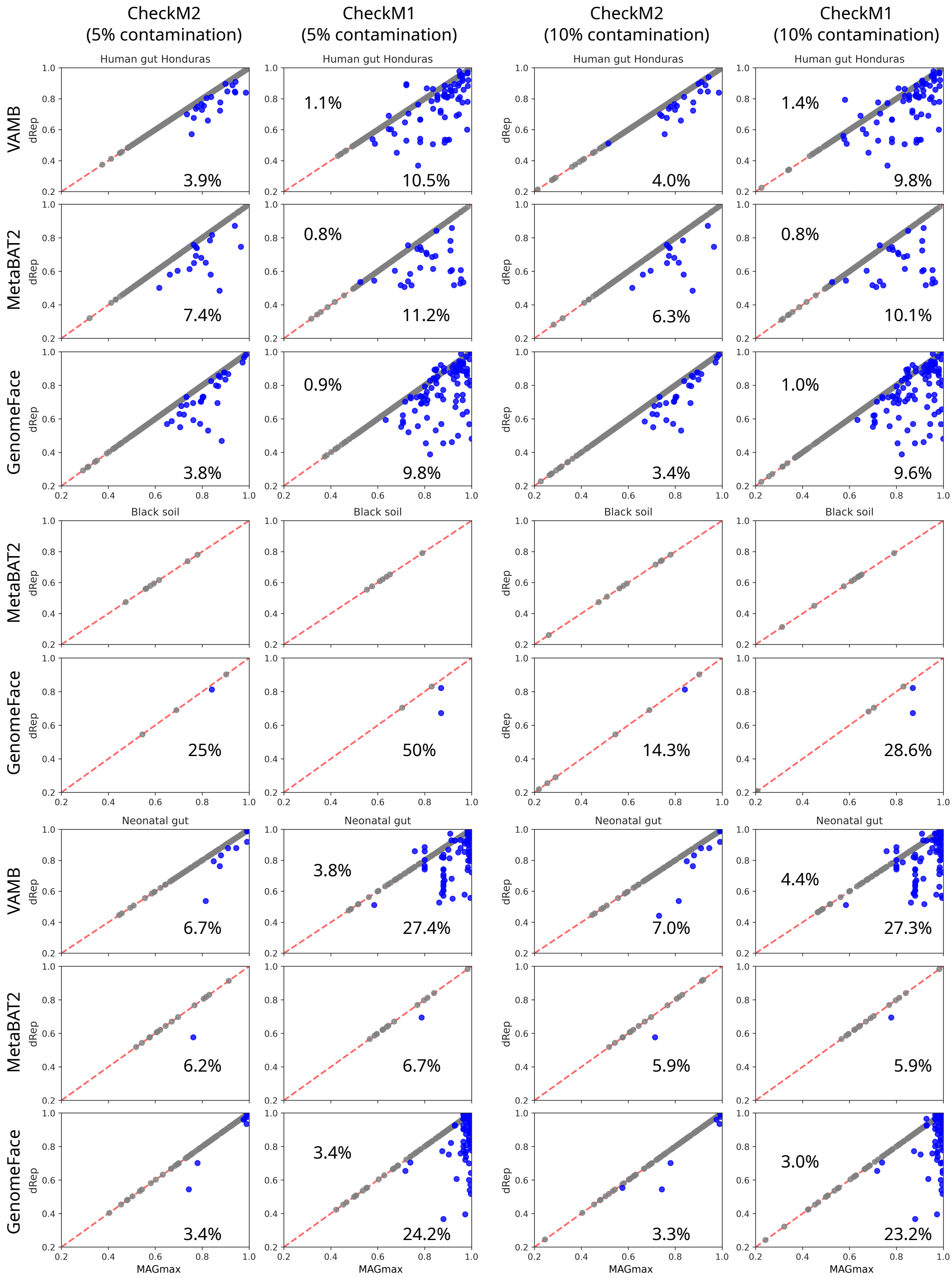


**Figure 2**: Comparison of quality scores of dRep bins and their corresponding MAGmax bins based on genomic cluster membership using CheckM2 (left) and CheckM1 (right) prediction. Numbers in percentage indicate the fraction of bins that differ and have a higher quality score than their counterpart in the other tool.

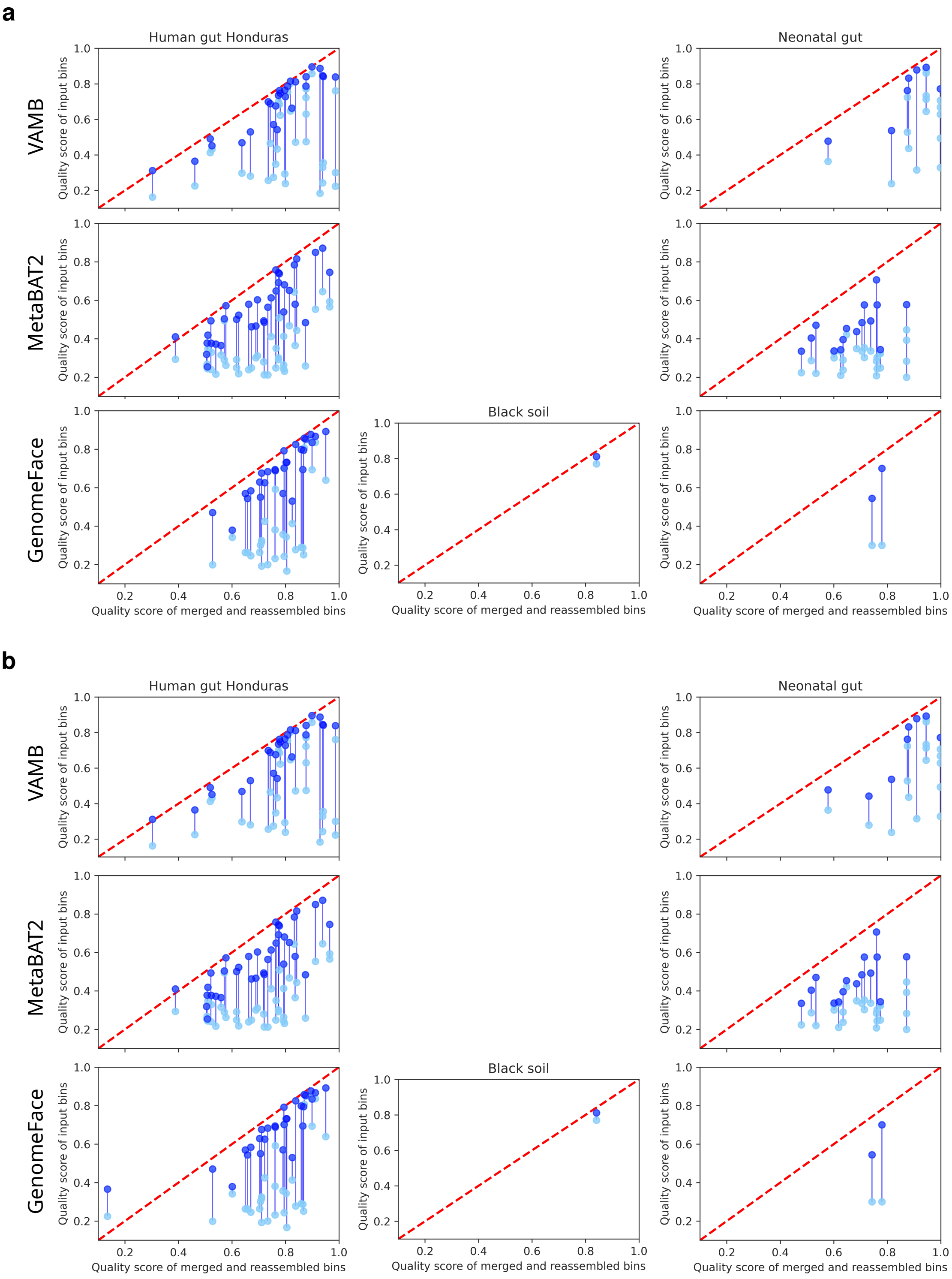


**Figure 3**: Comparison of quality scores between input bins (y-axis) and their corresponding merged and reassembled bins (x-axis) at 5% contamination (a) and 10% contamination (b). Blue circles represent the highest-quality bins within each genomic cluster, while cyan circles indicate the other input bins. Vertical lines connect the input bins that were used for merging and reassembly.
